# From lab to ocean: bridging swimming energetics and wild movements to understand red drum (*Sciaenops ocellatus*) behavior in a tidal estuary

**DOI:** 10.64898/2026.04.03.716345

**Authors:** BJ. Gibbs, JA. Strother, CR. Morgan, D. Pinton, A. Canestrelli, JC. Liao

## Abstract

Understanding how fish navigate complex natural environments requires bridging fine-scale biomechanics with ecological behavior. We investigated the volitional movement and energetics of wild red drum (*Sciaenops ocellatus*) across laboratory, mesocosm, and field settings. Using flow-respirometry, we quantified metabolic costs and swimming kinematics under ecologically relevant flow conditions shaped by bluff bodies mimicking mangrove roots and oyster mounds. Fish swimming in turbulent wakes exhibited reduced oxygen consumption and altered tailbeat dynamics, especially at high flow speeds. In a large outdoor mesocosm, dual accelerometers revealed a rich behavioral repertoire, including maneuvering and rest, which is not easily observable in confined lab settings. Spectral analysis and clustering identified eight distinct locomotory states, highlighting the limitations of summed acceleration metrics. Field telemetry tracked wild red drum across a 54 km estuarine corridor for a three-year period through an array of 36 acoustic receivers, revealing movement patterns shaped by tidal flow and physical habitats. Hydrodynamic modeling revealed that while laboratory trials demonstrated substantial energetic savings at high flows (approaching 100 cm/s), wild fish were detected predominantly in low-velocity microhabitats (<30 cm/s) near structurally complex features. This mismatch suggests that habitat selection is an adaptive strategy driven by ecological factors such as foraging opportunities, predation refuge, and site fidelity, rather than hydrodynamic efficiency alone. Our multi-scalar approach demonstrates that while flow-structure interactions can reduce locomotor costs for fish, habitat use in the wild reflects broader ecological constraints, offering a framework for integrating biomechanics, physiology, and ecology in conservation-relevant contexts.

## Introduction

The environment plays a foundational role in driving real-time animal behavior, in addition to sculpting morphology and physiology at evolutionary time scales (Lejeune et al., 1998; Dickinson et al., 2000; Full et al., 2000; Li et al., 2013). While laboratory studies have been instrumental in furthering our understanding of locomotion, especially under steady-state conditions, they are conducted within confined spaces that inherently constrain natural movement and behavior. Consequently, our understanding of how animals navigate in natural, complex environments remains limited. Work in terrestrial systems reveal that animals running over uneven terrain drastically change limb performance compared to running over smooth surfaces (Ferris et al., 1999; Daley and Biewener, 2006; Druelle et al., 2019). Laboratory studies often examine performance limits rather than investigate verified, ecological behaviors. For example, studies often focus on their remarkable running ability of ostriches (Abourachid and Renous, 2000; Rubenson et al., 2004), but their locomotion is characterized by slow walking when freed to behave in natural habitats (Daley et al., 2016). Studying how animals move in natural environments promises to reveal new insights into the energetics and ecology of locomotion.

Animals living in fluids must constantly contend with external forces from a three-dimensional medium. Dynamic flow conditions arising from the interaction of current with abiotic structures can shape locomotion and behavior for both invertebrates and vertebrates (Chabot et al., 2007; Cote and Webb, 2015; Mechalec et al., 2017). In the lab, naturalistic, unsteady flows have been shown to alter swimming behavior and energetics (Webb, 1998; Liao et al., 2003a; Fish and Lauder, 2006), but we know less about the mechanisms of how fishes behave in nature. Marine fishes in coastal habitats also experience daily fluctuations in the tidal cycle that generate large changes in current velocity (Pethick, 1980; Beck and Legault, 2012). These flows interact with submerged structures like mangroves, rocks and oyster reefs. These fluid-structure interactions increase hydraulic roughness and flow resistance (Wright et al., 1990; Styles, 2015) and alter local flow fields that fishes must navigate. Consequently, fluid–structure interactions not only modify the physical flow environment but also generate a heterogeneous energetic landscape (Shepard et al., 2013), imposing variable locomotor costs on fishes. Thus, linking physiology and movement kinematics to local hydrodynamic environments remains essential for understanding how fishes navigate and exploit dynamic energetic landscapes in natural coastal systems.

To understand the volitional movement and energetics of wild fish across different environmental contexts, we adopted an integrative experimental approach. This included controlled laboratory settings in a flow tank to measure swimming energetics, recording volitional behavior with accelerometers in large mesocosms, and monitoring individual fish movements in the wild where flow and habitat information were available. We selected the red drum (*Sciaenops ocellatus*) as a local species to investigate movement within natural environments, given their ubiquitous presence in local estuarine habitats in Florida and across the southern Atlantic Ocean (Matlock, 1987; Adams and Tremain, 2000). Using a variable speed flow-respirometry system we dissected the influence of flow with ecological structures abstracted from local natural habitats (e.g., oyster mounds and mangrove roots) on metabolic expenditure and swimming kinematics. In a large outdoor mesocosm pool, we examined how these same fish behaved in a more ecologically representative setting. Here, natural light cycles, habitat structures, and the presence of local prey allowed us to observe a richer behavioral repertoire than could be elicited in a flow tank respirometer. The mesocosm acted as a behavioral bridge, linking reductionist laboratory insights to the complexity of field behavior. Lastly, we utilized acoustic field telemetry to map large-scale spatial and temporal movement patterns of fish across an estuarine-wide landscape. To investigate how hydrodynamics influence fish movement patterns, we coupled a high-resolution Delft3D-FLOW numerical model of the Guana Tolomato Matanzas (GTM) estuary with concurrent acoustic telemetry data from 2019–2021.

Our laboratory experiments demonstrated that red drum preferentially occupy high-velocity flow immediately downstream of objects, where vortical structure and velocity gradients permit station holding with reduced energetics despite elevated free-stream speeds. However, field telemetry revealed that detections were concentrated in lower-velocity regions of an estuary, including nearshore areas adjacent to mangroves and oyster reefs. This apparent discrepancy suggests that habitat selection reflects scale-dependent hydrodynamic structure: fine-scale flow refugia embedded within high-flow environments may confer energetic advantages in specific settings, whereas broader landscape-level occupancy in nature may favor regions where ambient velocities (and thus baseline locomotor costs) are reduced. Reconciling these scales requires explicitly testing whether laboratory-derived flow preferences translate to field-scale occupancy when fish experience complex tidal forcing and heterogeneous energetic landscapes. The null hypothesis is that habitat use is independent of flow velocity at both microhabitat and landscape scales, such that laboratory flow preferences do not predict field-scale spatial occupancy.

Together, our lab-to-ocean approach offers a multi-scalar view of locomotion: the flow tank quantifies energetic principles, the mesocosm reveals behavioral diversity, and the field uncovers ecological relevance (Fig. 1). We believe that this approach is vital to contextualizing the relevance of laboratory experiments and charts a framework for future integrative studies in animal behavior.

**Fig. 1.**
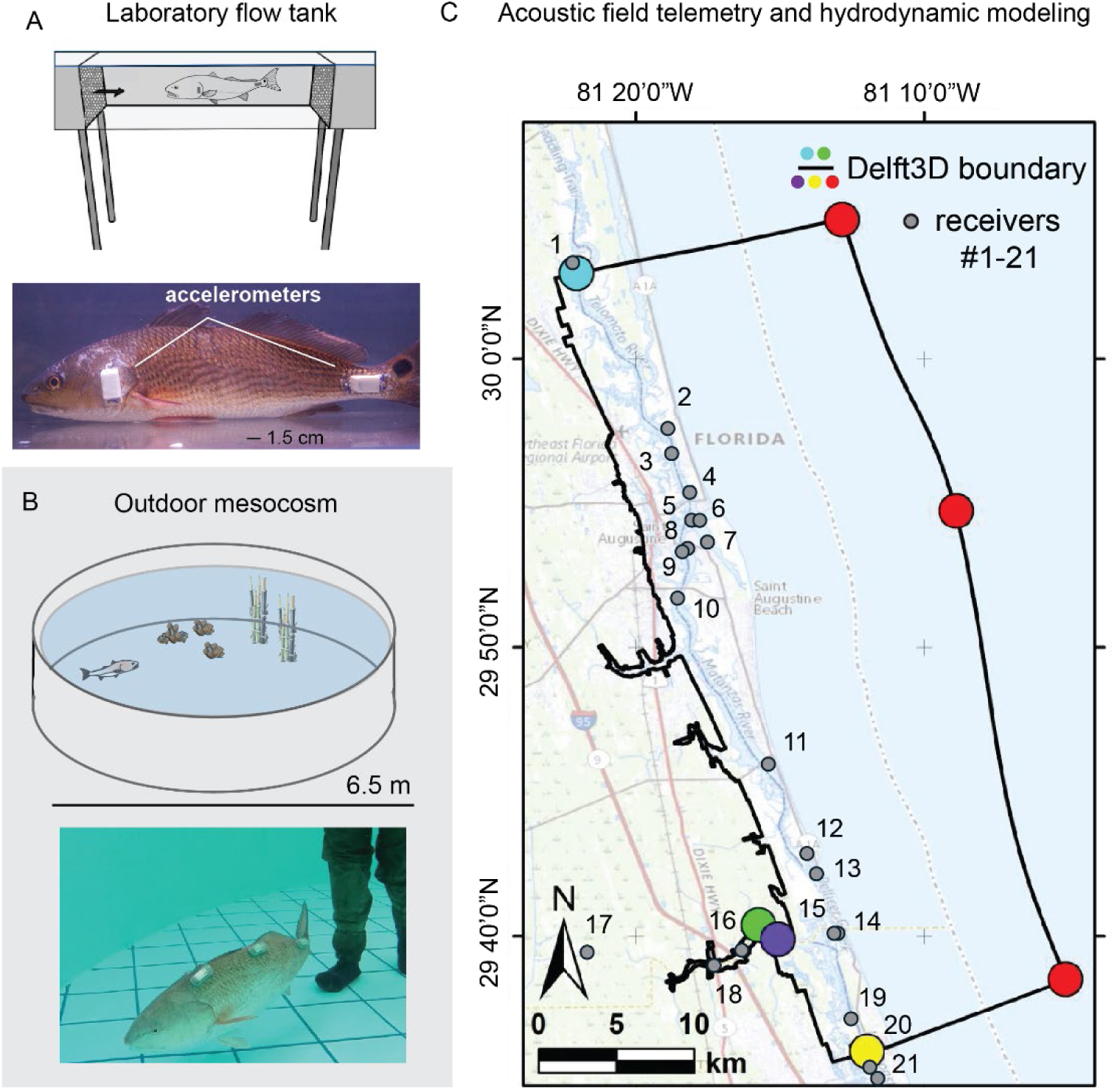
A lab-to-ocean approach to understand the volitional movement and energetics of wild red drum (*Sciaenops ocellatus*), illustrating three interconnected experimental scales used to bridge the gap between fine-scale biomechanics and large-scale ecological behavior. (A) By using a variable-speed flow tank, we measure metabolic rates and swimming kinematics in response to different flows and simplified habitat structures inspired by field observations. This setup also serves to calibrate accelerometer data, creating templates for interpreting behaviors in more natural environments. (B) A large outdoor mesocosm offers a realistic environment to observe behavioral repertoires under local photo periods, simulated habitat complexity, and natural prey. This intermediate scale of investigation bridges the lab and field, and when extended over long time periods, encourages a diversity of spontaneous behaviors such as maneuvering, feeding and resting. (C) Tracking wild fish over large spatial and temporal scales with acoustic telemetry, when integrated with hydrodynamic flow models, bridges fine-scale laboratory insights with habitat-dependent spatial behavior and ecological function. The map shows the Guana Tolomato Matanzas estuary, where the black line is the region of simulated flow (e.g. the Delft3D model domain boundary) where fish were acoustically tracked for three years. The colored circles are the boundary conditions of the model and the gray dots are the locations of the acoustic receivers, numbered progressively from north to south.

## Methods

### Animal care

For the flow tank and mesocosm experiments, wild red drum (n=18; 40.1 ± 4.2 cm mean ± standard deviation) were line-caught in the Matanzas River and housed in large (10000 L) indoor tanks with a flow-through seawater system at the Whitney Laboratory for Marine Bioscience. Fish were acclimated for at least three days before experimentation and were fed live shrimp daily. All experimental protocols were approved by the University of Florida IACUC (#202200000056).

For the field tracking experiments, wild red drum (n = 5; 75.3 ± 8.5 cm) were line-caught in the St. Augustine inlet area. Once onboard the boat, they were immediately placed in a holding trough with recirculating seawater. A local anesthetic (Vetericyn plus antimicrobial hydrogel; Innovacyn) and iodine (10% povidone-iodine) were applied to a small area on the fish’s belly anterior to the pelvic fins. A scalpel (Size 10; Cincinnati Surgical Company, Cincinnati, OH) was used to make a small 2-inch incision in this area, and an acoustic tag (V9 transmitter, ping-rate 60s-180s, Vemco, Boston, MA; 69 kHz) was magnetically activated and then implanted into the fish. The incision was sutured (2-0 gauge polydioxanone suture; Ethicon Inc., Somerville, NJ, USA), and antimicrobial cream (Vetericyn hydrogel; Innovacyn) was applied to the wound while the fish recovered. The fish were then released, to be detected whenever a tagged fish was within ∼300 m of a receiver. Since red drum is a popular recreational species that is kept by anglers for consumption, only fish classified as over-slot (27 inches or larger; > 68.5 cm) were selected for tagging to reduce the risk of a tagged fish being harvested.

### Kinematic analysis

High-speed video (100 fps, Phantom Miro 340; Vision Research, Wayne, NJ, USA) was captured to determine swimming kinematics under three hydrodynamic treatments inspired by estuarine habitats: (1) uniform free-stream flow (control), (2) holding station behind a 2D bluff body (5 cm D-shaped cylinder simulating a mangrove root), and (3) holding station behind a 3D bluff body (10 cm × 10 cm spherical mound simulating an oyster cluster) (Fig. 2). Video recordings of 20 tailbeats or more were randomly captured during swimming bouts in each treatment across flow velocities found in the local estuarine environment: 22 cm/s, 61 cm/s, and 100 cm/s. A swimming bout was defined as active swimming characterized by lateral body movements within the region directly downstream of each bluff body. Videos were analyzed using DeepLabCut (Mathis et al., 2018) to label 8 points on the body midline (e.g., snout, operculum, pectoral fin base, pelvic fin base, anal fin bases, caudal fin base, caudal fin tip). Midline coordinates were then processed using NumPy. Coordinate locations were tracked over the course of a tailbeat so that the pixel distance over time could be used to derive three kinematic variables: tailbeat amplitude, tailbeat frequency, and head angle. Tailbeat amplitude was calculated as the distance between the caudal fin tip at the onset and conclusion of a half tailbeat cycle, while tailbeat frequency was calculated as the number of full tailbeat cycles in 60 s periods throughout the entire swimming duration. Head angle was calculated using the center of mass (pectoral fin base) and the snout at the onset and conclusion of a half tailbeat cycle.

**Fig. 2.**
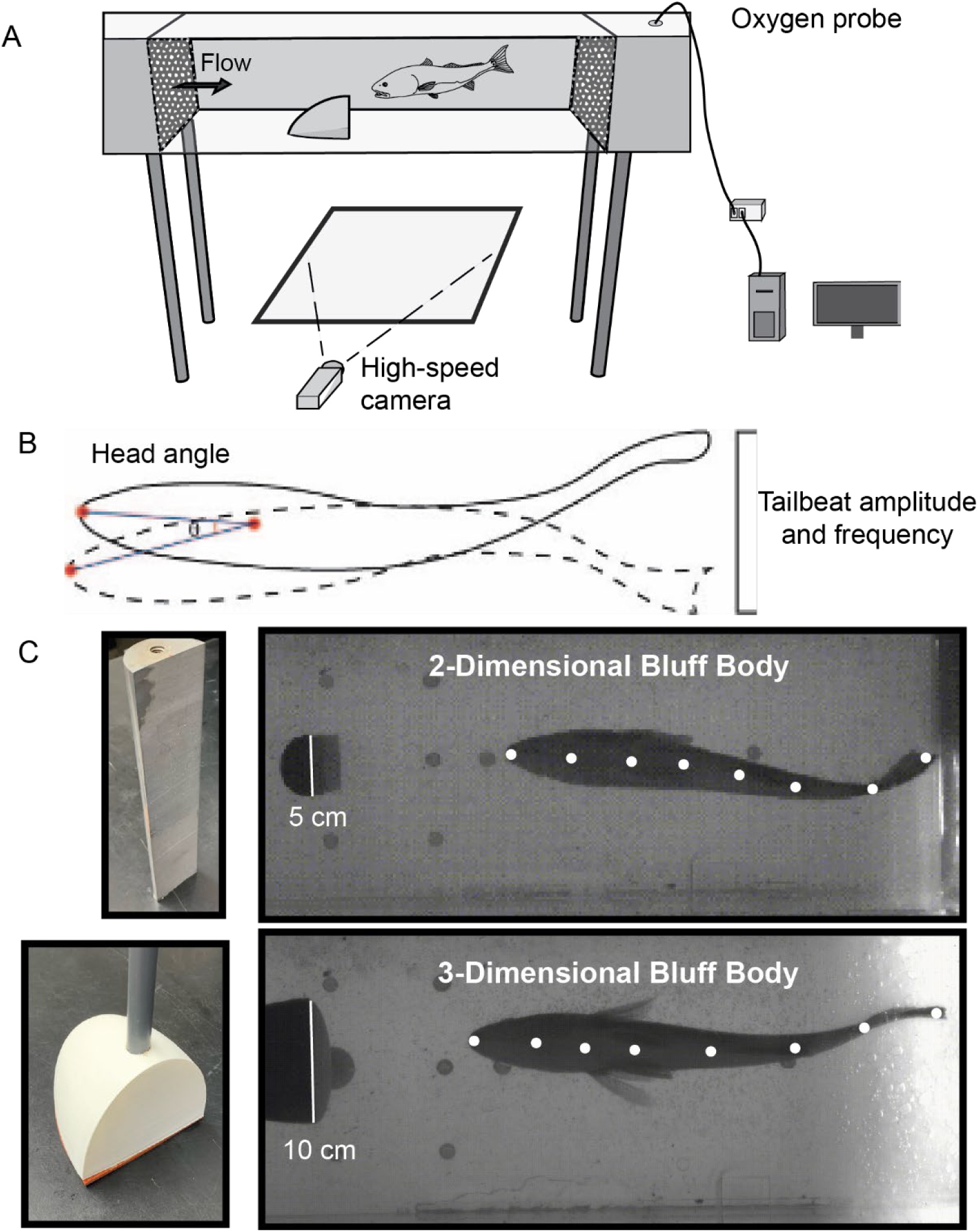
Experimental set up to measure oxygen consumption and kinematics of wild red drum. (A) Schematic of the flow tank respirometer used to swim animals at three ecologically-relevant flow speeds. (B) Three kinematic variables were measured simultaneously with respirometry experiments: head angle, tailbeat amplitude and tailbeat frequency. (C) Experimental treatments included a 2-dimensional and 3-dimensional bluff body used to simulate natural structuresin the environment.

### Respirometry

Oxygen consumption of red drum was measured concurrently with kinematic recordings using the same three hydrodynamic treatments described above. Measurements were conducted in the 185 L flume recirculating flow tank with an oxygen probe (Fig. 2) and monitored with Autoresp v.1.6 software (Loligo Systems, Denmark). Prior to experimentation, a background level of oxygen consumption was measured to account for any microbial life in the tank.

Individual red drum (n = 5; 37.7 ± 1.3 cm) were allowed to acclimate overnight in the flume running at low speed to establish a continuous swimming rhythm. After the acclimation period, a trial was started in which active swimming behaviors were observed and oxygen consumption was measured for three 15-minute measurement cycles. For every oxygen measurement cycle, a 300 s flush, 60 s equilibration, and 900 s measurement period sequence was applied following the intermittent flow respirometry methodology (as described by Johansen et al., 2020). Oxygen consumption data were averaged and plotted using NumPy and Matplotlib in Python. A two-way ANOVA was used to assess the effects of treatment, swimming speed, and their interaction on swimming kinematics and oxygen consumption. Specific significant effects between treatments and speeds were further analyzed using Tukey’s post-hoc tests.

### Volitional behaviors in a mesocosm

Wild red drum (n=13; 41.3 ± 3.1 cm) were anesthetized in a 10 L tank (20 ± 1 ^◦^C) containing tricaine methanesulfonate (45 mg/L MS-222; Finquel Inc., Argent Chemical Laboratories Inc., Redmond, WA, USA) buffered with sodium bicarbonate (90 mg/L NaHCO3). Once a fish was unresponsive to a tail pinch with forceps, it was transferred to a surgery trough (10 L) with recirculating seawater. A three-axis accelerometer data logger (Axy5 S, Technosmart Europe, Rome, Italy) was sutured (2-0·gauge polydioxanone suture; Ethicon Inc., Somerville, NJ, USA) to the fish’s operculum and second data logger was sutured to the tail base. After surgery the fish was then transferred to larger, circular tank with flow-through seawater (750L) and allowed to recover overnight. Immediately after the recovery period, the accelerometers were turned on which recorded acceleration and timestamp (UTC) data continuously (50 kSPS) for the 5-day experiment.

To understand how rectilinear swimming speed is encoded by our accelerometers, each accelerometer on the swimming fish was calibrated in the 185L flow tank in uniform flow. Once the desired flow speed was set, swimming kinematics were recorded for 90 seconds. The trial was then terminated to avoid fatiguing the animals. Each swimming bout was captured with high-speed video (Phantom Miro Lab 340, 100 fps, Vision Research, Wayne, NJ, USA) and stored in files with timestamps (UTC). After the swimming bout was completed, the accelerometers were physically tapped to produce a spike in the accelerometry data (e.g. to provide a time point reference). Manual video review determined the time interval of steady swimming following the acceleration spike. This interval was then identified in the accelerometry data by locating the tap-induced spike as a temporal reference point and verifying timestamp alignment between video and accelerometer records. This period was then used to calculate tailbeat frequency and amplitude for the selected flow tank speed, as described below. After these calibration experiments, the fish was transferred to a circular tank with flow-through seawater (750 L) and allowed to recover overnight. After 24 hours the fish was transferred to a 7,000 gallon (diameter = 6.5 m, height = 0.8 m) outdoor mesocosm pool for 5 days. The mesocosm was populated with naturalistic habitat structures and native prey (e.g. white shrimp *Litopenaeus setiferus* and gulf killifish *Fundulus grandis*).

Each accelerometer was calibrated by orienting the housing so that the x, y, or z axis aligned with the gravitational vector. An affine transform was then used to convert the raw acceleration vector to an acceleration vector aligned to the housing and with the correct scale (1g). Since the tail and opercular accelerometers were not intrinsically synchronized it was necessary to perform a post-hoc analysis to align the recorded data. After the accelerometers were attached to the animal, each anesthetized animal was briefly rotated such that the gravitational acceleration produced a distinct signature in the measured acceleration values. Cross-correlation of the acceleration during this interval was used to identify the start time delay between the accelerometers. Clock drift over the experimental period was removed using an analogous procedure. It was assumed that the acceleration vector in the dorso-ventral direction was similar for both the tail and opercular accelerometers, the cross-correlation of this component of the acceleration was computed for each 2 h block and used to identify the delay, and times were then offset and linearly stretched to align data from the two sensors. The accelerations were then band-pass filtered to remove low-frequency drift and high-frequency noise (0.2 - 20 Hz, 4th-order Butterworth), and the major axes of variation were identified through a principal components analysis (PCA). The first principal component appeared to represent tail-beating or ventilatory cycles and was used for subsequent analysis. For each interval, the movement frequency was calculated by computing the Fourier Transform of the acceleration along the first principal axis, identifying the frequency with the greatest signal power, and using peak interpolation to determine the precise central frequency. Similarly, the movement amplitude was computed as the root-mean-square (RMS) of the acceleration along the first principal axis. We examined the relationship between the swimming speed dictated by the tank flow speed and each computed accelerometry metric (tailbeat amplitude, tailbeat frequency, opercular amplitude, opercular frequency). The curves were then fit to the following empirically selected function

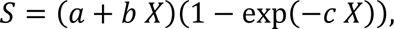

where S is the speed, X is the examined metric, and a, b, and c are constants. The tailbeat amplitude provided the norm of residuals and was used when subsequently estimating swimming speed from accelerometry data.

Accelerometer data for the mesocosm were analyzed similarly to the flow tank experiments. Data was analyzed using custom-written MATLAB scripts (Mathworks R2023a). The recorded time was aligned to the zeitgeber time (ZT) by offsetting the recorded time values (UTC) by the sunrise time, which was calculated with the NOAA sunrise/sunset calculator (https://gml.noaa.gov/grad/solcalc/sunrise.html) using the date of each experiment and the GPS coordinates of the mesocosm. After experiments were completed, the accelerometers were removed and fish were released back into the wild.

To understand the different swimming behaviors exhibited by the animals we analyzed the accelerometry data using dimensionality reduction and clustering. As described above, acceleration vectors were pre-processed by 1) transforming them into a standardized coordinate system defined by a PCA analysis and then 2) performing low-pass filtering. The data was then divided into blocks (8s duration), the data was subsampled by 4x to reduce computational requirements, and then the frequency spectrum was calculated for each block using a discrete Fourier transform. This procedure generated a dataset composed of 3-dimensional spectra for both the tail and respiratory acceleration vectors from all recorded individual animals. This dataset was then processed using t-SNE non-linear dimensionality reduction to project the data points onto a 2-dimensional plane. The 2-dimensional points were then analyzed using k-medoids clustering in order to group instances into 8 classes. A representative spectrum was then produced for each class by taking the median across all data points within the group.

### Acoustic Telemetry

To understand behavior in the field, we positioned an array of 21 acoustic receivers (VR2W, Innovasea, Boston, MA) along the GTM National Estuarine Research Reserve (GTMNERR), spanning 54 km from Matanzas Inlet to St. Augustine Inlet. Receivers were attached to 8-foot aluminum poles that were bolted into dock pilings above the low tide water line, with the permission from private citizens, park organizations, and the US Coast Guard. Securing the posts during low tide enabled a dry-deployment system, allowing a team of two researchers to pilot a boat to retrieve, download data and redeploy receivers. Note that receivers were often fixed to docks which were adjacent to oyster flats and mangroves. Collaborations with the FACT Network, Animal Telemetry Network, Georgia Department of Natural Resources and the Ocean Tracking Network (SECOORA) provided additional access to over 5,000 receivers from the Gulf of Mexico to Nova Scotia, facilitating the effective tracking of both inshore and offshore movements.

Movement data was downloaded during routine receiver checks and processed in the VUE software. This data was utilized to compare movement patterns of detected individuals based on location, time of year, time of day, and tidal cycle. R scripts were used to plot fish movement and provide summary statistics.

### Hydrodynamic modeling

We employed the Delft3D-FLOW model (Deltares 2025) to simulate the hydrodynamics of the GTM estuary, resolving unsteady flow driven by tidal and meteorological forcing on a regular, boundary-fitted grid. The model domain (outlined in black in Figure 1c) covers the GTMNERR and is centered near St. Augustine, Florida, USA. Grid resolution varies from approximately 30 m × 100 m in the offshore region to about 15 m × 20 m within the GTM estuary. Ocean bathymetry was obtained from NOAA’s bathymetric inventory, while estuarine bathymetry integrates data from multiple sources, including the Florida Natural Areas Inventory vegetation map and topographic and bathymetric LiDAR datasets from USGS, USACE, and NOAA. The model was originally developed in Pinton and Canestrelli (2020), calibrated in Gray et al. (2021), and has been successfully applied to hydrodynamic simulations for Pinton and Canestrelli (2024). Simulations were conducted from 1 January 2019 through 31 December 2021, corresponding to the period during which acoustic receivers were deployed to track fish movements in the GTM estuary. The Delft3D-FLOW model was run with a computational time step of 1 min. Boundary forcing included: (i) harmonic constituents of the astronomical tide derived from three nearby NOAA tide stations (Fig. 1C, red circles); (ii) observed water levels from GTMNERR and Florida Department of Environmental Protection stations located at the northern and southern ends of the Intracoastal Waterway (blue and yellow circles); (iii) tidally filtered freshwater discharge from the USGS Pellicer Creek gauging station (green circle); and (iv) meteorological forcing, including wind speed, wind direction, and precipitation, measured at the Pellicer Creek meteorological station (purple circle).

To investigate the hydrodynamic conditions experienced by tagged fish in the GTM estuary, we utilized the known coordinates of 21 acoustic receivers (gray dots in Figure 1C, numbered sequentially from north to south) deployed throughout the estuary. Using output from the Delft3D-FLOW hydrodynamic model, we extracted horizontal velocity magnitude and direction within a 50-meter radius around each receiver location. These velocities were retrieved continuously over the entire simulation period (January 1, 2019, to December 31, 2021) at a 30-minute output interval, and specifically at the time instances when detections for five individuals (i.e., Fish 23332, 53427, 23326, 59223, and 59225) were recorded by the receivers.

## Results

### Swimming kinematics and oxygen consumption in flow

We measured tail beat frequency, tail beat amplitude, and head angle to assess swimming kinematics under the flow conditions simulated by our model and confirmed with measured current velocities obtained from the Army Corps of Engineers (Fig. 3A-C). Two-way ANOVA revealed that both treatment and flow speed significantly affected tailbeat amplitude and tailbeat frequency as there was a significant interaction between our flow treatments and flow speed for both metrics (Tailbeat amplitude: F = 25.25, p < 0.001; Tailbeat frequency: F = 39.67, p < 0.001). Post-hoc Tukey comparisons showed specific differences between treatment-speed combinations. For example, tailbeat amplitudes for 2D bluff body treatments generally differed significantly from both the free stream and 3D bluff body treatments at all speeds, whereas 3D bluff body and free stream treatments did not differ significantly at intermediate speeds, such as 61 cm/s (p = 0.95). Additionally, two-way ANOVA of head angle showed significant main effects of treatment (F = 32.62, p < 0.001) and speed (F = 207.23, p < 0.001) but no significant interaction (F = 0.73, p = 0.58), suggesting that hydrodynamic treatments influenced head orientation consistently across all flow speeds.

**Fig. 3.**
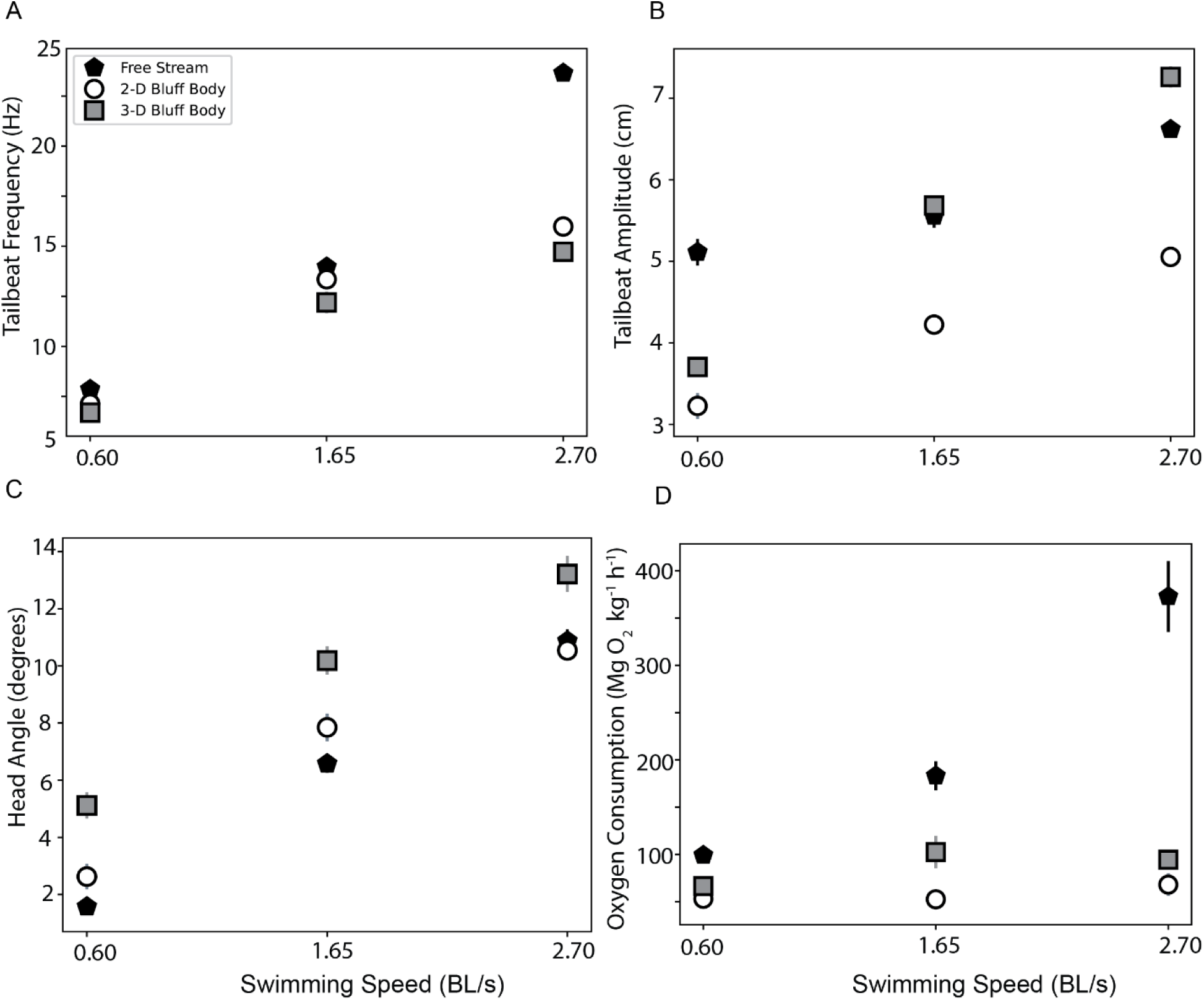
Kinematics and oxygen consumption of red drum under different hydrodynamic conditions. (A) Tailbeat frequency (Hz) of the three treatments plotted against swimming speed. At the highest swimming speeds, the two turbulent flow treatments displayed much a lower tailbeat frequency than in uniform flow. (B) Tailbeat amplitude (cm) of the three treatments plotted against swimming speed. Tailbeat amplitude was lowest while swimming behind at 2-D bluff body at all speeds while highest while swimming behind the 3-D bluff body at the two highest swimming speeds. (C) Head angle (degrees) of the three treatments plotted against swimming speed. Across all swimming speeds, head angle was the greatest while swimming behind a 3-D bluff body. (D) Oxygen consumption of the three treatments plotted against swimming speed. At all swimming speeds, oxygen consumption was lower while swimming behind a bluff body compared to uniform flow. Error bars represent std. dev. and may be obscured by symbols.

We measured oxygen consumption (MO₂) to assess the energetics of swimming under different bluff body treatments and flow speeds (Fig. 3D). Two-way ANOVA revealed that both bluff body treatments and flow speeds significantly affected MO₂ consumption with a significant interaction between bluff body treatment and speed (MO₂: Treatment: F= 64.42, p < 0.0001; Speed: F= 31.36, p < 0.0001; Treatment + Speed: F= 18.88, p < 0.0001). Post-hoc comparisons indicated that MO₂ consumption increased sharply with swimming speed in free stream treatments, whereas 2D bluff body treatment showed relatively little change across speeds, and 3D bluff body treatment displayed intermediate increases.

### Behavior in the mesocosm

Calibration of accelerometers on fish in a flow tank revealed that both tailbeat and opercular acceleration and frequency increased with swimming speed (Fig. 4A-D). Lines of best fit were used to correlate accelerometer movement with natural swimming patterns found when fish were allowed to behave freely in the mesocosm. We found that fish in the mesocosm exhibit different behaviors than fish swimming steadily in the flow tank, and generally preferred to swim at lower speeds (Fig. 5A-B). Behavior also varied between individual animals, with some animals demonstrating slow locomotion over extended periods of time while other animals exhibited bouts of slow and fasting swimming. Spectral analysis and t-SNE dimensionality reduction analysis suggested that swimming behaviors fall within a broad continuum rather than distinct groups (Fig. 6A). To understand the variation within this continuum, the behavioral space was divided into 8 clusters, and we computed the median frequency spectra for each cluster. Based on a qualitative interpretation of these spectra, the clusters were labeled as Swimming 1-4 (representing increasing speeds), Sedentary, Breathing, Slow Maneuvering, and Fast Maneuvering. Swimming 1-4 behaviors were characterized by tail and respiratory accelerations with similar frequency spectra. In both, the majority of the signal power was concentrated within the first principal axis of variation in a narrow peak, at a frequency that increased from Swimming 1 to 4. This is representative of simple sinusoidal movements. Sedentary behavior exhibited low signal power for both tail and respiratory acceleration, while Breathing behavior showed high signal power for respiratory acceleration but low power for tail acceleration. The Slow and Fast Maneuvering behaviors exhibited moderated signal power for both tail and respiratory acceleration, but with spectra that differed from those in the Swimming 1-4 behaviors. For both maneuvering behaviors, the signal power was spread across multiple principal axes and a broad range of frequencies. This is likely representative of complex, non-repeating movements.

**Fig. 4.**
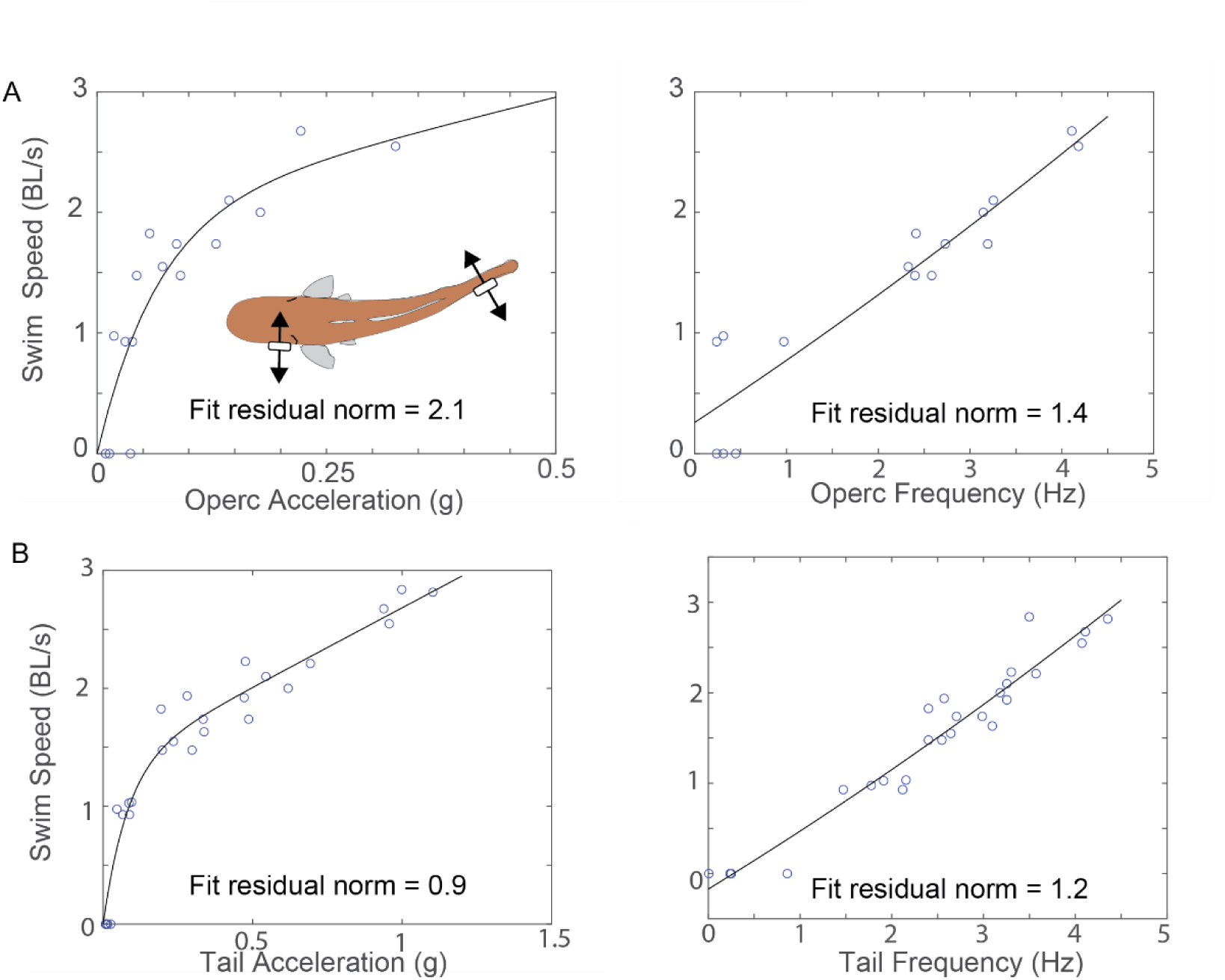
Dual accelerometer calibration of red drum in a flow tank flume. (A) Tri-axial accelerometers were surgically attached to a fish’s operculum and tail base were subject to a range of known flow speeds in our flow tank flume to correlate accelerometer movement with natural swimming patterns. (B) Opercular acceleration and frequency plotted against swimming speed. (C) Tail acceleration and frequency plotted against swimming speed. Lines of best fit were used to determine the most reliable metrics to correlate accelerometer movement with natural swimming patterns.

**Fig. 5.**
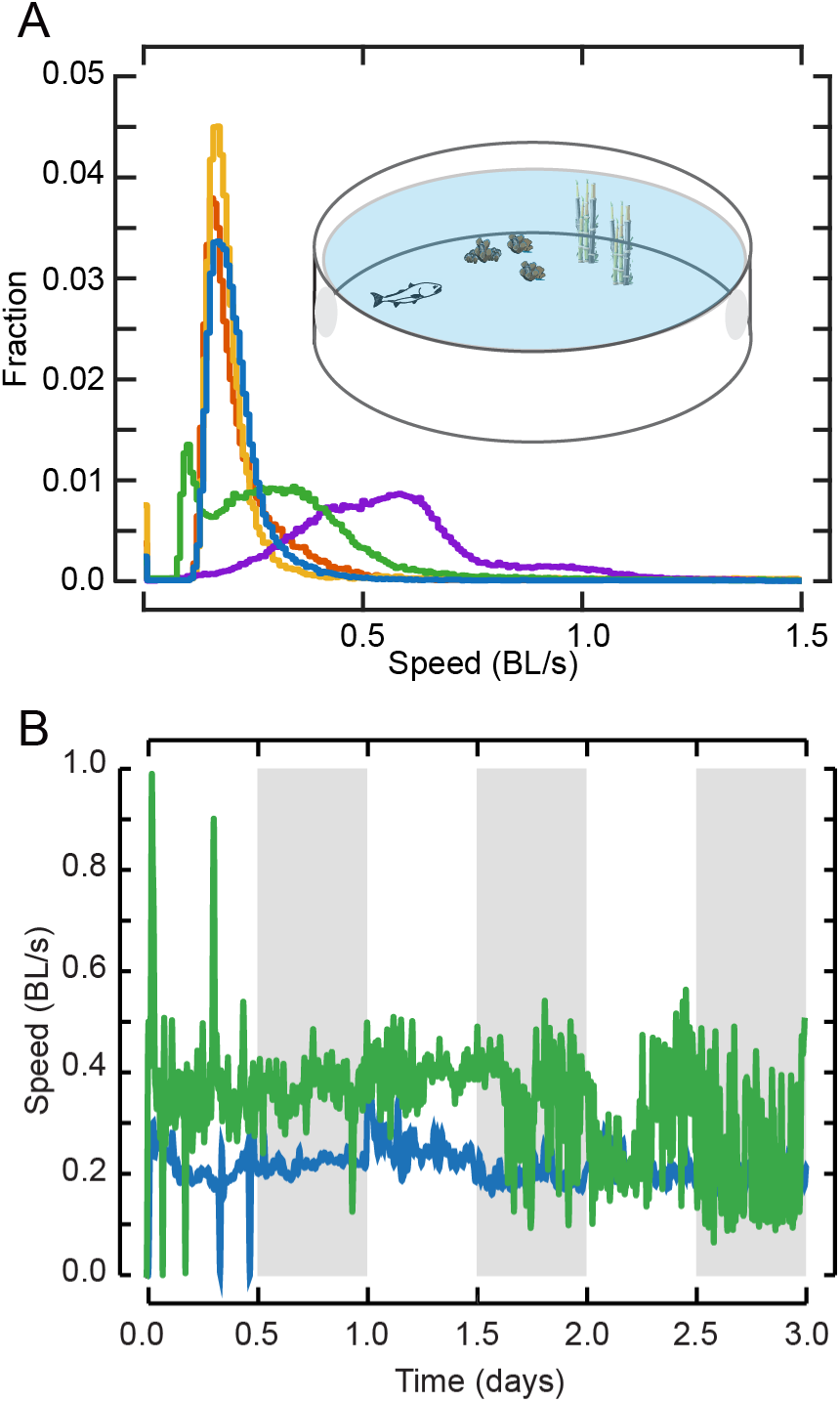
Swimming speed of red drum in a large outdoor mesocosm. (A) Histograms of swimming speed for individual animals in mesocosm (n=5, estimated from tail acceleration, see Fig. 4). (B) Time series of swimming speed for two representative animals locomoting within the mesocosm (individuals are annotated with the same color in panels A&B). The time series begins with sunrise on the first experimental day, and grey bars indicate approximate periods of darkness (approx. 12h:12h light:dark during trial days).

**Fig. 6.**
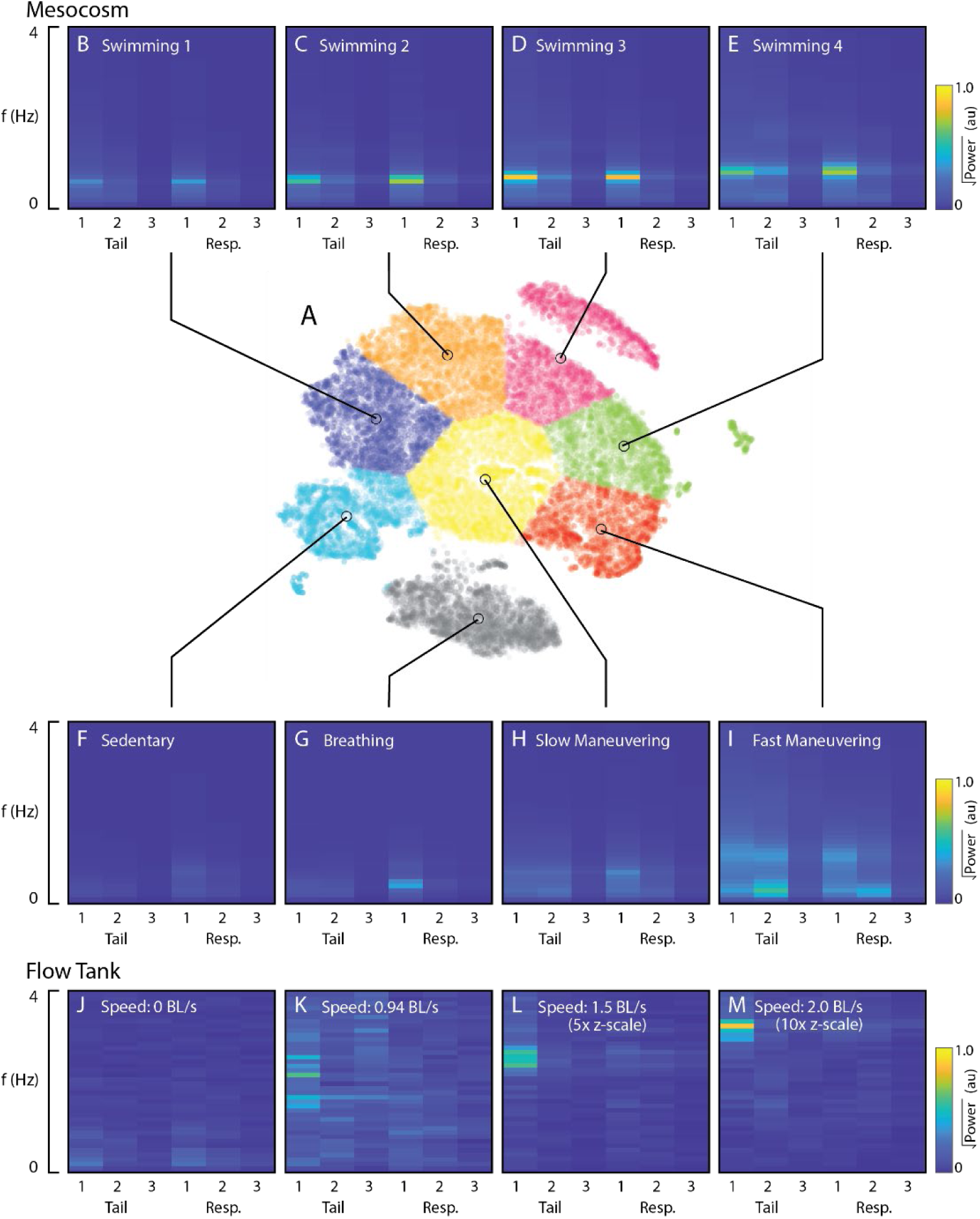
Redfish display diverse swimming behaviors in mesocosm and flow tank environments. (A) Low dimensional representation of different swimming behaviors exhibited by redfish in the mesocosm, generated by performing a t-SNE analysis on spectra of tail and respiratory accelerations. (B-I) Median spectra for each cluster within A. Spectra are shown as pseudo-color graphs with warmer colors depicting greater values for signal power. The y-axis represents different frequencies, and the x-axis shows the 3 standardized coordinates of the tail and respiratory acceleration vectors. (J-M) Similar to B-I except showing the spectra for swimming behaviors displayed in the flow tank at different flow speeds.

The behaviors observed in the mesocosm were compared to those exhibited within the flow tank experiments by computing the frequency spectra for the tail and respiratory acceleration (Fig. 6B). We found that the signal power for these behaviors was concentrated within the first principal axis of tail acceleration. At low swim speeds the spectrum was broad, while the spectrum narrowed to a narrow peak with high amplitude as the swimming speed increased. Comparison of the mesocosm and flow tank datasets suggests similarity between the mesocosm Slow Maneuvering behavior and the flow tank 0 body lengths per second (BL/s) behavior, and similarity between the mesocosm Fast Maneuvering behavior and the flow tank 0.94 BL/s behavior. However, overall our data suggest that animals show markedly different swimming behaviors in a large mesocosm compared to those observed in a flow tank.

### Movement and flow patterns in the wild

Acoustic telemetry revealed that fish traversed a diverse mosaic of coastal habitats, including mangrove shorelines, oyster reefs, mud flats, and oceanic inlets (Fig. 7A). For example, an individual animal (fish 23332) moved∼14 km across the St. Augustine inlet region, accumulating thousands of detections along the inshore waterway from November 2019 to November 2020 (Fig. 7B). In January 2020, this fish moved ∼9 km offshore for six days before returning to the inlet, revealing the extent that red drum movement patterns can span multiple habitat types and hydrodynamic regimes.

**Fig. 7.**
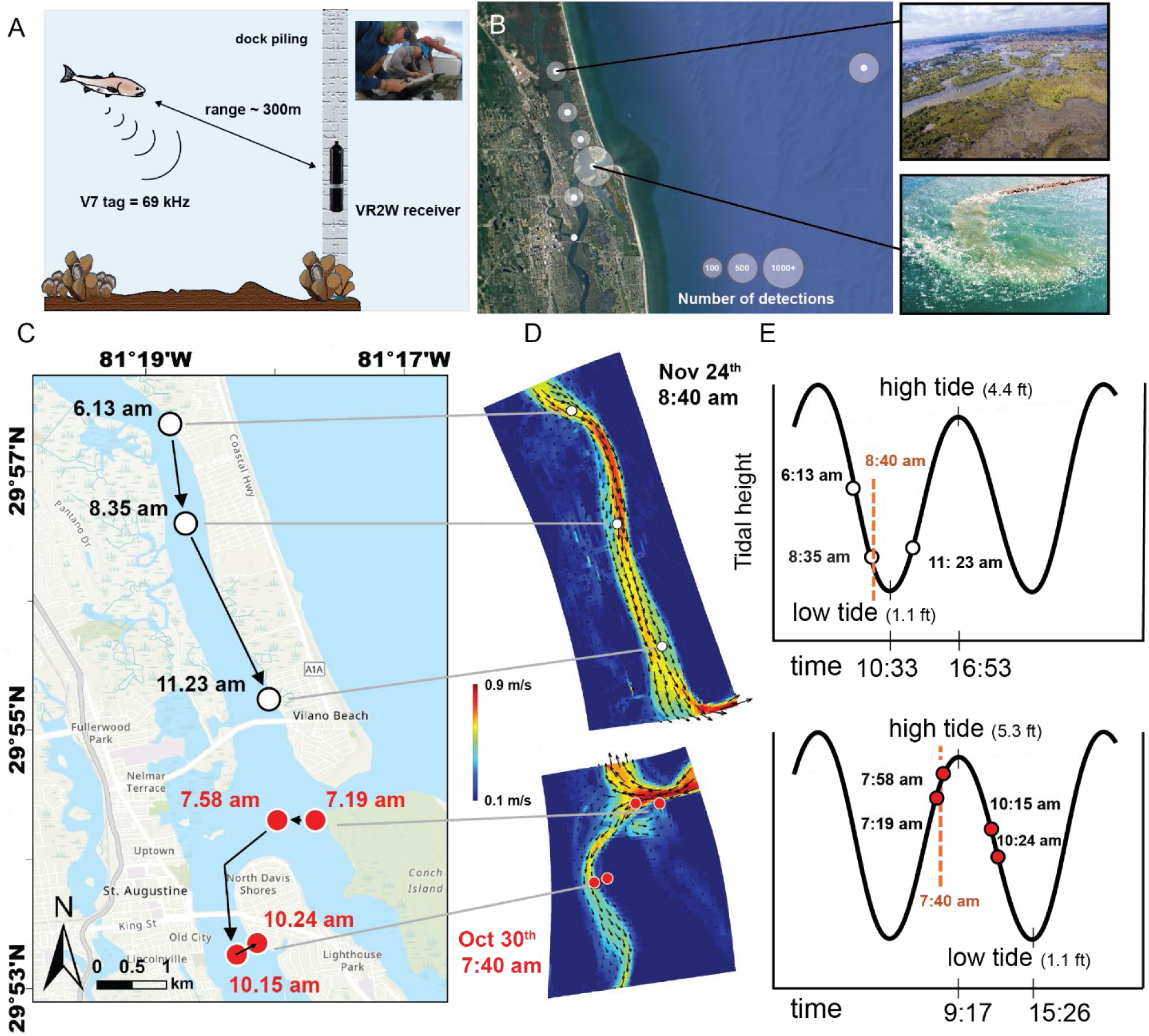
Red drum movement in a tidal estuary. (A) Schematic illustrating how red drum movements are monitored with stationary receivers after acoustic tags are surgically implanted into individual fish (inset picture). (B) Movement patterns of an individual fish as it moved through diverse inshore and offshore habitats. White dots indicate receiver location and grey circles denote number of detections. (C) Temporal tracking data of a fish on two different dates: October 30^th^ and November 24^th^ 2020. (D) Delft3D computational simulations of these regions on this day reveal the magnitude of the average velocity experienced by this fish. (E) The time of fish detection at specific receivers indicates that fish move with and against the flow during the tidal cycle.

Fish move both with and against tidal flows. For example, another animal (fish 53427) moved ∼4 km south from the St. Augustine inlet (Oct. 30^th^ 2020, red circled receivers), swimming both with (detection at 7:58 am) and against (detection at 10:15 and 10:43 am) moderate to low tidal currents (incoming high tide at 9:17 am, Fig. 7C-E). The same fish was detected November 24^th^, 2020 near the Guana River moving south (white circled receivers) with the outgoing tidal current ∼4.5 km toward the inlet mouth (detection at 6:13 and 8:35 am, low tide at 10:33 am), as well as with the incoming tide (detection at 11:23 am). These movement patterns during times when the simulated average flow velocities ranged from 0.1–0.9 m/s.

A broader analysis of individuals in the field compared the flow velocities coincident with fish detections at receivers with the flow velocities across the entire numerical simulation period (Fig. 8A, blue and orange bars, respectively). These results indicate that fish detections are biased toward relatively low flow velocities near the receivers (blue bars). Nearly 90% of detections occur at velocities below 0.50 m/s, with the majority (∼75%) concentrated between 0.15 m/s and 0.40 m/s. By comparison, velocities below 0.50 m/s and within the 0.15–0.40 m/s range occur for approximately 80% and 45% of the total simulated period, respectively, over the three years of analysis in the vicinity of the receivers (orange bars). This mismatch suggests that fish are preferentially detected near receivers, most of which are adjacent to the shoreline, during periods of low to moderate flow, particularly when velocities range between 0.15 m/s and 0.40 m/s.

**Fig. 8.**
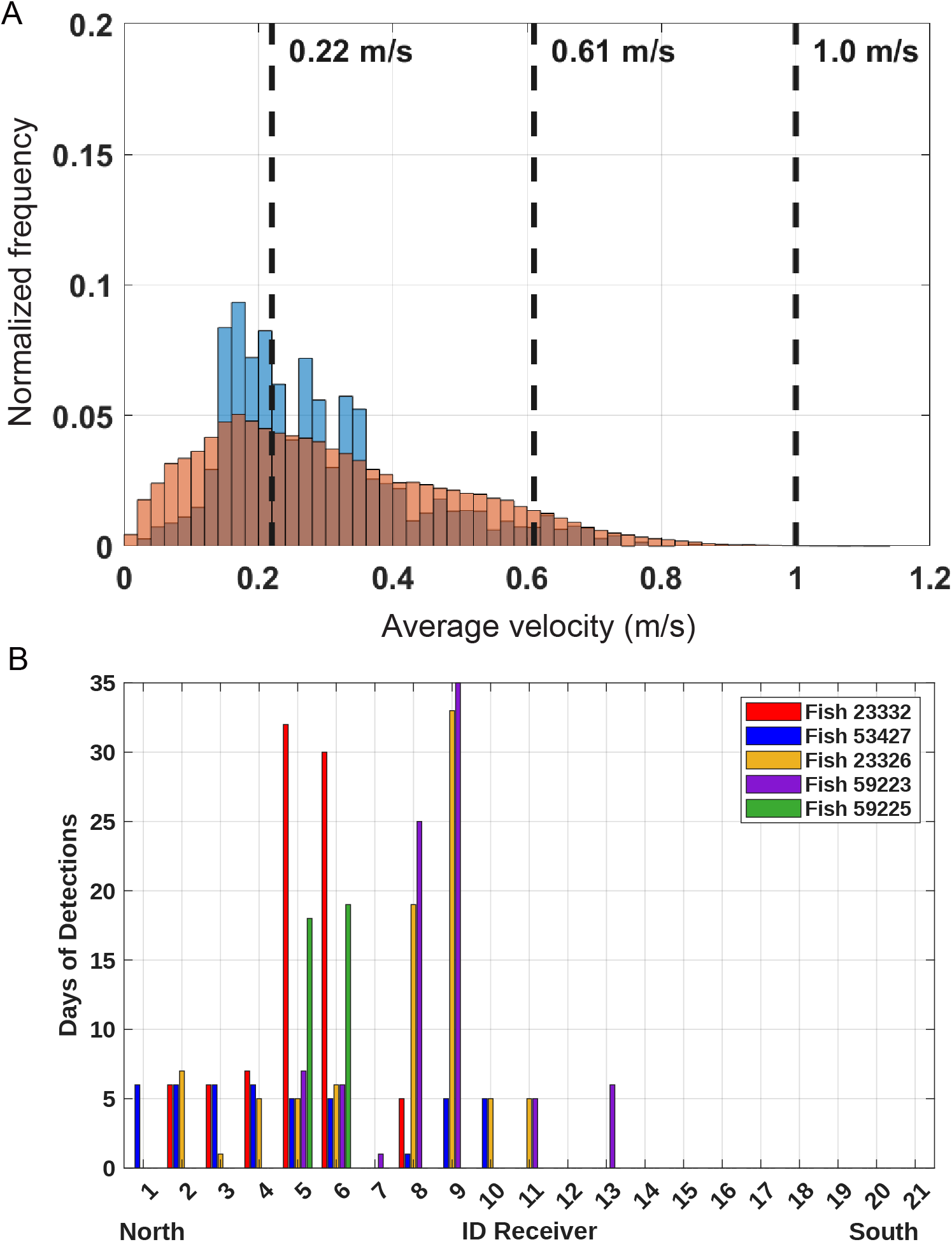
(A) Distribution of velocity magnitudes at the receivers for five individual fish. The blue bars indicate velocities recorded at times when fish detections occurred. The orange bars represent the overall velocity distribution throughout the entire Delft3D-FLOW simulation. (B) Spatial distribution of detections for all five fish. Each bar represents the number of days a fish was detected at a given receiver.

The number of days each fish was detected at a given receiver (Fig. 8B) reveal that certain individuals (fish No. 23332 and 59225) remain near specific receivers near the St. Augustine Inlet (receivers No. 5 and 6), suggesting a strong site preference. Similarly, fish No. 23326 and 59223 predominantly remained near receivers located midway between St. Augustine Inlet and the confluence of the Matanzas River and the San Sebastian River (receivers No. 8 and 9). Notably, only these fish were detected for a limited number of days in the southern portion of the estuary (i.e., the watershed influenced by the Matanzas Inlet and monitored by receivers from 11 to 21), indicating their tendency to remain within the zone of influence of St. Augustine Inlet. However, certain individuals (fish No. 53427) were detected at receivers located farther inside the GTM estuary (receivers No. 1 to 4 and 8-10), indicating a broader spatial distribution and the absence of a preferential region.

## Discussion

### Energetic and Behavioral Strategies Across Scales

A comprehensive understanding of animal movement is challenging given the difficulty in reconciling fine-scale kinematic data with behaviors expressed over ecologically relevant spatiotemporal scales. Expanding understanding of the biomechanical, physiological and neural mechanisms underlying natural movement depends on technologies and approaches that enable new measurements and analyses (Nathan et al., 2008, Yartsev and Ulanovsky, 2013; Mao et al., 2020; Das et al., 2023).

Here, we advocate for an integrated approach to fish locomotion, where laboratory experiments are inspired by and occur alongside behavioral measurements in the field (Fig. 1). Mesocosms represent an intermediate step, offering environmental complexity to promote natural behaviors while retaining experimental control. This multi-scaled approach provides a cohesive framework for understanding how different scales of study inform one another. In recent decades, the advent of biologging technologies have provided key tools for researchers trying to understand the movements of wild animals *in situ*. In birds and bats, these devices can collect and record a wide range of information, such as the animal’s satellite-derived location and environmental parameters, providing long-term datasets that elucidate large-scale migrations and habitat use patterns (Jouventin and Weimerskirch, 1990; Richter and Cumming, 2008). For marine taxa, however, the aquatic medium imposes substantial limitations on data transmission due to impedance differences between water and air, hindering wireless data transmission and requiring tag detachment and retrieval to recover data (e.g. delayed data retrieval of tags that surface upon release). Pop-off satellite archival tags have met these challenges for large pelagic species and revealed movement and congregations to unique marine habitats such as offshore seamounts and submarine canyons (Block et al., 2001; Sedberry and Loefer, 2001; Carlson et al., 2010). Typically equipped with low frequency accelerometers, these tags can monitor static and dynamic accelerations across multiple axes and offer insights into the locomotion of freely behaving marine animals (Cavagna et al., 1961; Dubois et al., 1976). While static acceleration can reveal changes in body posture, dynamic acceleration can reveal overall body accelerations (Shepard et al., 2010; Papastamatiou et al., 2021). Placed on the dorsal fin or body trunk, these devices can infer gliding behaviors and rapid accelerations indicative of prey pursuit (Holland et al., 2009; Nakamura et al., 2011; Brown et al., 2013). However, accelerometers do not provide the temporal or spatial resolution that biomechanists operate on when seeking to understand locomotory mechanics in the lab (e.g. millisecond/millimeter scale). While laboratory approaches enable precise kinematic and metabolic measurements under controlled conditions and simplified environments, they are often limited by spatial scale and behavioral diversity.

These reductionist approaches are essential but incomplete without validation in natural settings where broader behavioral repertoires are expressed (Drucker, 1996, Dewar and Graham, 1994a; Muller et al., 2008). Insights into locomotion are best revealed when observing animals in large, natural spaces (Golet et al., 2006; Gleiss et al., 2019). For example, fishes tracked in their natural habitat display behaviors not observed in the laboratory, revealing ecological drivers of locomotion (Nakamura et al., 2011; Papastamatiou et al., 2021, Ellerby and Gerry, 2011; Jones et al., 2007; Kendall et al., 2007; Ellerby et al., 2018).

Our results lead us to suggest that timing windows occur when the energetic cost of locomotion can be reduced under specific situations when flow and habitat interact. To explore this hypothesis mechanistically, we replicated field-relevant flow conditions in a laboratory setting, using both two-dimensional and three-dimensional obstacles to simulate natural topographies found in the inshore habitat of red drum (e.g. mangrove roots, dock pilings, rocks and oyster reefs). In nature, fish routinely interact with structures and flow (Eggertsen et al., 2016; Laurioux et al., 2024; Liao, 2025) and may reduce energetic costs (Taguchi and Liao, 2011, Roche et al., 2014, Liao et al., 2003a; Fulton et al., 2013; Zhang and Lauder, 2024). Our respirometry trials revealed a reduction in oxygen consumption and altered swimming kinematics when individuals swam in the wake of these structures (Fig. 3D), with the magnitude of energetic savings influenced by obstacle geometry. Specifically, fish swimming behind two-dimensional objects exhibited lower metabolic rates than those swimming behind three-dimensional objects, despite equivalent flow velocities. This may reflect the reduced demands of postural stabilization in two-dimensional wakes, which typically exhibit more predictable vortex shedding patterns (Liao et al., 2003b; Taguchi and Liao, 2011). These differences were greatest at higher flow speeds, which are available naturally to red drum throughout the GTM NERR estuary (Fig. 7D). Observing red drum in the field together with laboratory energetic measurements indicates that structure–flow interactions can enhance swimming efficiency, and that use of structurally complex habitats during tidal transitions likely reflects adaptive responses to hydrodynamic variation.

Our accelerometry data revealed substantial intra- and inter-individual variability in swimming speed throughout the day (5A-B), consistent the finding that animals rarely exhibit uniform velocity in naturalistic settings (Fuiman and Webb, 1988; Daley et al., 2016). Fishes, like terrestrial animals, spend substantial time maneuvering and resting. Variations in swimming speed occurred among individuals despite inhabiting the same environment. This variability may reflect underlying individual traits, linked to social status or personalities, which can modulate locomotor performance and decision-making (Krause et al., 2000; Nakayama et al., 2016; Webster, 2017). While a single trunk-mounted accelerometer can capture gross body movement, it remains insufficient for disentangling overlapping behaviors and must be interpreted cautiously. For instance, opercular acceleration may increase in response to elevated metabolic demand during sustained swimming, but it can also be elevated during stress responses, complicating interpretation (Smit, 1965; Webb, 1971). Additionally, a single accelerometer may produce inaccurate readings by failing to account for transient translational accelerations caused by incidental movement or contact with the external environment. To address these limitations, we employed a dual-sensor configuration, with one accelerometer affixed to the caudal peduncle and another to the operculum. This arrangement enabled us to separate propulsive and ventilatory signals, yielding higher behavioral resolution. Correlation of body kinematics to known swimming velocities in a flow tank allowed us to derive acceleration templates corresponding to specific behavioral states (Fig. 6). Similar approaches have been validated in both freshwater and marine species and have advanced our ability to reveal the full diversity of behaviors that animals exhibit in the wild (Getz and Kline, 2019; White and Lauder 2025). For example, calibrating accelerometers with visual observations of swimming in a mesocosm allowed nocturnal foraging rates in bonefish (*Albula vulpes*) to be revealed in the wild (Brownscomb et al., 2014). Importantly, we found that summating dynamic accelerations from a single 3-axis accelerometer, a common approach in field studies (Thiem et al., 2015; Ward et al., 2019; Lopez et al., 2022), may mask functionally distinct behaviors. Our results suggest that analysis of the frequency spectra of each axis enables a wide variety of behaviors to be distinguished. For example, the “Swimming” and “Fast Maneuvering” behaviors have similar summed accelerations, while analysis of the spectra indicates that these are clearly distinct behaviors. The “Swimming” behaviors exhibit tail accelerations along a single axis at a fixed frequency, while the “Fast Maneuvering” behaviors exhibit tail accelerations along multiple axes with broad spectra reflecting irregular motions. These findings underscore the need to integrate spectral characteristics of acceleration data when inferring behavior. In future work, multiple-accelerometer approaches to understanding unconfined teleost behaviors will take place in the field, where programmed tag detachment has enabled data recovery (White and Lauder 2025).

When calibrated appropriately, accelerometry provides a powerful bridge between the laboratory and the field, enabling researchers to quantify behaviorally relevant variation in energy use and movement dynamics. A substantial gap exists in our understanding of the sensory inputs and processing that drives the motor behaviors of freely-swimming fishes (Musall et al., 2019; Rynes et al., 2021). Future studies may benefit from coupling accelerometry with complementary tools such as magnetometers, video tracking, and neural sensors. For example, recent work using neurologgers has enabled real-time monitoring of sensory processing and motor commands in freely moving fishes (Vinepinsky et al., 2017; Gibbs et al., 2023), offering new opportunities to link environmental perception with behavioral output. By coupling accelerometers to neurologgers we can gain a deeper understanding of ecological behaviors. For example, simultaneous recording of body accelerometers with lateral line afferent activity could elucidate how flow sensing, modulated by afferent and efferent sensory gain, contributes to locomotor decisions during predator-prey interactions or schooling (Montgomery and Bodznick, 1994; Lunsford et al., 2019; Skandalis et al., 2021). Mesocosms cannot fully replicate natural complexity but remain a critical intermediate step toward ecologically valid experimentation.

Hydrodynamic modeling reveals that the estuary where our red drum were collected spans a wide velocity range, with peak flows approaching 100 cm s⁻¹ in channels but substantially lower velocities near shorelines where acoustic receivers were deployed (Fig. 1). Tagged fish detections were strongly biased toward the lower end of this velocity spectrum (<30 cm s⁻¹; Fig. 8), despite the availability of much faster flows elsewhere in the system. Importantly, receivers were mounted on nearshore docks adjacent to oyster reefs and mangrove shorelines, habitats that provide structural complexity independent of their hydrodynamic effects. While our laboratory respirometry approach demonstrates substantial energetic savings behind oyster- and mangrove-mimicking structures at high flow speeds, fish in the wild were rarely detected at receivers during such conditions. This mismatch suggests that fish are not occupying these habitats primarily to exploit hydrodynamic refugia during peak flows, as our lab studies would suggest. Instead, association with structures adjacent to our receivers at low velocities likely reflects non-hydrodynamic drivers, such as foraging opportunities, refuge from predation, spatial memory, or site fidelity to benthic features. Similar decoupling between energetic optima and habitat use has been suggested in other fishes (Moustaka et al. 2024) where structurally complex habitats are selected for ecological function rather than locomotor efficiency alone. Together, these results indicate that while flow–structure interactions can reduce energetic costs under extreme conditions, the day-to-day habitat choices of wild red drum appear to be governed by broader ecological constraints, with hydrodynamic benefits emerging opportunistically rather than dictating routine space use. These findings therefore lead us to reject the null hypothesis that habitat use is independent of flow velocity at both microhabitat and landscape scales. However, the direction of the relationship is not predicted by laboratory experiments alone: fine-scale flow preferences measured under controlled conditions do not translate directly to landscape-level occupancy, underscoring that energetic optima and ecological habitat choice may operate through distinct mechanisms across scales.

One aspect of the present study concerns the spatial resolution of acoustic telemetry within the GTM estuary. Although current velocities in the estuary can be relatively high during certain tidal phases, the precise swimming paths of tagged fish within this channel remain unknown. Acoustic receivers have a detection radius of approximately 300 m, a detection therefore confirms fish presence within a broad area rather than pinpointing their position within the cross-sectional flow profile. Fish may preferentially use lower-energy microhabitats within or adjacent to the estuary. Examples include salt marshes, mangroves and oyster reefs fringing the inlet, marina basins, and wetland edges flanking the southern and northern receivers where flow velocities are typically below ∼0.5 m/s. If fish routinely select these refugia, the hydrodynamic conditions they experience could differ from the channel-averaged velocities extracted at receiver locations. This uncertainty could be addressed by examining the sensitivity of the results to receiver placement and detection radius, and by integrating acoustic detections with finer-scale positioning methods to better resolve individual movement paths relative to local hydrodynamic conditions. Resolving this uncertainty is critical to testing the scale-dependent hydrodynamic hypothesis. If fish consistently occupy low-velocity microhabitats within otherwise high-flow regions, it would suggest that the energetic principles revealed in the laboratory do operate in the field, but at a spatial grain finer than current telemetry can resolve.

Our results contribute to a growing body of work advocating for integrative approaches that merge biomechanical, physiological, and ecological perspectives on locomotion. Our findings raise several points of integration with broader hydrodynamic ecology frameworks.

First, the intersection of fine-scale flow heterogeneity and behavioral positioning underscores the importance of microhabitat structure in shaping individual energy budgets and movement ecology. Second, laboratory-measured energetic optima do not necessarily predict natural behaviors. The apparent discrepancy between energy savings quantified with respirometry at high flow velocities and actual field distributions partially supports the scale-dependent hydrodynamic hypothesis. Fish appear to exploit flow refugia opportunistically at fine scales while selecting habitats at the landscape scale based on broader ecological drivers. This highlights a nuanced relationship between flow and habitat use, where energetic principles established in the laboratory remain valid but must be considered within the context of broader ecological constraints operating at larger scales. Future respirometry experiments could align more closely with lower flows reflective of conditions that fish encounter naturally. Third, the role of sensory systems in detecting and exploiting such hydrodynamic refugia warrants exploration, as positional adjustments directly influence behavioral decisions in spatially variable flows. Finally, these results have implications for habitat restoration and conservation design: preserving or restoring structurally complex features such as oyster reefs and mangrove habitats may enhance populations of estuarine fishes by maintaining conditions that encourage growth and survival. By employing an integrative approach that considers insights from lab and field, we can begin to unveil the locomotory diversity and behavioral strategies that enable fish to thrive in diverse and changing habitats.

## Acknowledgements

We would like to thank Esra Gokturk, Glen Greenwald, Peter Meyers, John Perkner, Luke Blaser and Old City Fly Shop for helping with preliminary analyses, tagging fish and maintaining acoustic receivers. We would like to acknowledge our funding support NSF (IOS 1856237, IOS 1932707, PHY 2102891, NSF CMMI 2345913), The Whitney Lab Mathei Scholars Fund, Patagonia Action Works, Yamaha Right Waters and Skeeter boats.

